# Beige adipose tissue-derived extracellular vesicles: a potent metabolic regulator and a novel remedy for nonalcoholic fatty liver disease

**DOI:** 10.1101/2024.01.01.573808

**Authors:** Kai Zhang, Sha Zhang, Bing-Dong Sui, Yuan Yuan, Lu Liu, Si-Qi Ying, Cheng-Han Li, Kai-Chao Zhang, Shu-Juan Xing, Zhi-Wei Yang, Yang Sun, Li-Juan Yu, Jin Liu, Yan Jin, Chen-Xi Zheng, Liang Kong

## Abstract

Adipose tissue (AT) is an essential metabolic and endocrine organ, which are categorized into the white adipose tissue (WAT) and the brown adipose tissue (BAT), as well as the beige adipose tissue (BeAT) that derives from WAT browning. Notably, extracellular vesicles (EVs), especially tissue-produced EVs, have been recognized to be critical players in multiple physiological and pathophysiological settings and provide efficient strategy for disease treatment. However, little is known about the BeAT-derived EVs (BeEVs). In this study, we induced BeEV formation by cold stimulation and extracted EVs from three kinds of AT *via* enzyme digestion combined with gradient centrifugation. The isolated BeEVs possess typical morphological, structural and marker characteristics of EVs. Proteomics analysis further demonstrated that the BeEVs hold a distinct protein profile while are enriched with multiple metabolic regulatory proteins. Importantly, systemic injection of BeEVs into mice improved metabolic dysfunction in a nonalcoholic fatty liver disease (NAFLD) model induced by high-fat diet (HFD). Moreover, we demonstrated that the BeEVs alleviated liver steatosis *via* decreasing the lipogenesis. These findings promote the understanding of AT-EVs and shed light on the isolation, identification and applications of BeEVs as a potent metabolic regulator and novel remedy for metabolic diseases.

## 1 Introduction

Adipose tissue (AT) is an important metabolic and endocrine organ, which actively responds to environmental cues and contributes to metabolic homeostasis maintenance, with coordinative functional regulation of multiple organs, such as liver, muscle, brain, and heart [1–3]. Recent studies have revealed that extracellular vesicles (EVs), a type of phospholipid bi-layered biological nanoparticles that enable intercellular bioactive molecule transfer and signaling transduction, play a pivotal role in the crosstalk between local cell types in AT and other organs [4–7]. While most studies have focused on EVs derived from cell culture supernatants or body fluids, EVs isolated directly from tissues have been gradually recognized to be critical players in multiple physiological and pathophysiological settings, which possess the features of parent cells and can comprehensively reflect tissue specificity and microenvironment [8–10]. Also, due to its easy access and high yield, tissue-derived EVs are expected to be promising candidates in the therapeutic applications of various diseases [8, 9]. As for AT, despite a few pioneering studies [11–13], the features and functions of adipose tissue-derived EVs (AT-EVs) remain largely unknown, especially considering the heterogeneous and plasticity of AT.

AT was previously categorized into two major types, *i.e.* the white adipose tissue (WAT) that is the main depot storing metabolic energy in the form of triglycerides (TGs) and the brown adipose tissue (BAT) that contributes to energy expenditure with non-shivering thermogenesis [2]. Due to its thermogenic function, BAT has been revealed to be promising in treating obesity and other metabolic diseases [14, 15]. Notably, it has been reported that EVs derived from BAT (BEVs) can lower body weight and blood glucose, and restore hepatic and cardiac functions in obese mice [12]. However, given that BAT is nearly absent in adulthood, the clinical applicability of BEVs is extremely restricted. Intriguingly, within recent years, research have revealed that WAT can be browned and converted into a new kind of AT, namely beige adipose tissue (BeAT), under specific conditions, such as cold exposure, starvation, exercise, and hormones [16–19]. In specific, it has been revealed that BeAT possesses BAT-like features and functions while being equipped with the abundant content of WAT [20–22], thus offering a favorable alternative source for extracting EVs. For now, the isolation method, cargo identification, and therapeutic application of BeAT-derived EVs (BeEVs) have not yet been established.

Nonalcoholic fatty liver disease (NAFLD) has become a leading liver disease with an estimated global prevalence of 25% [23]. Of note, NAFLD is commonly associated with a cluster of metabolic dysfunctions, such as obesity, dyslipidemia, hypertension, and diabetes, but lack of effective treatments [24]. As two pivotal tissues in lipometabolism, the metabolic activities of the liver are often modulated by the AT, in which EVs seem to be closely involved [5, 13, 25, 26]. Also, it has been revealed that the liver is an ideal target for EV-based therapeutic strategies, due to the tropism of systemically injected EVs for it [27]. For example, our previous study has shown that mesenchymal stem cell (MSC)-derived EVs mainly target the liver after intravenous injection and alleviate high fat diet (HFD)-induced hepatic steatosis [28]. Nevertheless, whether BeEVs can target the liver *in vivo* and play a potential therapeutic role in NAFLD is unknown. Investigation of the biodistribution and the therapeutic effects of BeEVs will provide a promising strategy for the treatment of NAFLD.

In this study, we first induced BeAT formation *via* cold stimulus and successfully isolated EVs derived from the three kinds of AT with enzyme digestion followed by gradient centrifugation. Then, we characterized the morphological features of BeEVs as well as of BEVs and WEVs. Next, we identified the protein profiles of BeEVs *via* LC-MS/MS analysis, which showed that BeEVs hold a distinct protein profile and are enriched with multiple metabolic regulatory proteins. More importantly, *in vivo* experiments showed that exogenous transplantation of BeEVs significantly alleviate HFD-induced fatty liver *via* inhibiting the lipogenesis in mice. Taken together, our study has established a stable and accessible isolation method for BeEVs, identified their morphological features and protein cargos, and revealed their therapeutic efficacy for NAFLD, thus promoting the understanding of tissue EVs and the establishment of EV-based therapies for metabolic diseases.

## 2 Materials and Methods

### 2.1 Animals

Animal experiments were performed following the Guidelines of Institutional Animal Care and Use Committee of the Air Force Medical University and the ARRIVE guidelines (IACUC Issue No.: IACUC-2023-kq-059). Male C57BL/6J mice were purchased from the Laboratory Animal Center of the Air Force Medical University. Mice were housed under a specific pathogen-free environment (24□, 12 h light/dark cycles, and 50% humidity), and were kept feeding and drinking *ad libitum*.

For the induction of BeAT, cold exposure, one of the commonly used methods with easy operation and high efficiency, was used. Eight-week-old male C57BL/6J mice were placed in a cryogenic rearing unit which kept a low temperature of 4□ while the other conditions remain normal. After consecutive cold stimulation for 7 days, the mice were removed from the rearing unit and sacrificed for subsequent experiments. All manipulations for cold-stimulated mice and mice-derived tissue were performed at 4□ or on ice.

For the establishment of HFD-induced NAFLD model, four-week-old male C57BL/6J mice were placed on a HFD (60% kcal from fat; D12492, Research Diets, USA) or a normal chow diet until the end of the experimental protocol. After 4 weeks of HFD feeding, the mice were intravenously administrated with AT-EVs (appropriately 100 μg based on protein measurement) or equivalent amount of PBS once a week for 8 weeks. Then, the mice were sacrificed with euthanasia, and the tissues were sampled and analyzed.

### 2.2 Harvest of the brown, white, and beige AT

After sacrifice, the mice were immobilized on the operating table. For the harvest of BAT, incisions were made in the midline of the back to expose the subcutaneous tissue under the scapular region. After removal of fascia and surface WAT, BAT, which was symmetrically distributed, was carefully extracted [29]. For the harvest of white and beige AT, incisions were made in the abdomen to expose the inguinal region bilaterally, and the subcutaneous inguinal AT was carefully collected as WAT (from room temperature-housed mice) or BeAT (from cold-stimulated mice). To detect the validity of cold-induced BeAT formation, haematoxylin and eosin (H&E) staining, and immunofluorescent (IF) staining of BAT marker uncoupling protein 1 (UCP1) were performed on the harvested BeAT, as stated below.

### 2.3 Isolation of AT-EVs

AT-EVs were isolated according to our previous established protocol [30]. Briefly, 1 g fresh and minced AT was placed in 15 mL centrifuge tubes containing 10 mL Liberase enzyme solution (Sigma-Aldrich, USA, diluted at 1:1000) and incubated at 37□ for 30 min. Then, the samples were centrifuged at 800 g for 10 min to remove the cells in the pellet. The supernatant was collected and centrifuged at 2,000 g for 10 min to remove the cellular debris. After that, the supernatant was purified through a 450 nm filter, transferred to 1.5 mL EP tubes, and centrifuged at 16,000 g for 30 min at 4□. The pellet was washed twice with PBS and the resulting AT-EVs were suspended with relevant solutions. All AT-EVs were stored at 4□ and used for experiments within 24 h.

### 2.4 Morphologic identification of AT-EVs

Nanoparticle tracking analysis (NTA) was conducted by PMX ZetaView (Malvern, UK) equipped with an EV 520 nm model to evaluate the size distribution and particle number of AT-EVs. Fresh AT-EVs solution was diluted at 1:1,000-20,000 in PBS to achieve a final concentration of 1×10^8^ particle/mL. The performance of particle analysis and generation of analysis reports were conducted according to the manufacturer’s instructions.

Transmission electron microscope (TEM) observation was performed on a FEI Tecnai G2 Spirit Biotwin TEM (Thermo Fisher, USA) to capture the morphological features of AT-EVs. To prepare the samples, the AT-EV pellet was suspended in 2% paraformaldehyde (PFA), and 20 μL AT-EVs were transferred to 200-mesh copper grids coated with formvar and dried for 5 min at room temperature. After excess suspension was removed with filter paper, the AT-EVs were negatively stained with uranyl acetate (Sigma-Aldrich, USA) for 2 min at room temperature, washed with distilled water and dried. Samples were observed at operating voltage of 100 kV, and photos were captured with a PHURONA camera (EMSIS, Germany) and analyzed with RADIUS 2.0 software (EMSIS, Germany).

Cryo-electron microscopy (Cryo-EM) observation was performed on an FEI Tecnai G2 instrument operating at 200 kV (Thermo Fisher, USA). To prepare the samples, the AT-EV pellet was suspended in 2% PFD, and 20 μL AT-EV were transferred to Quantifoil R2/1 holey carbon grids and plunge-frozen into liquid ethane with an FEI vitrobot Mark IV set at 4°C and 95% humidity. Vitrified grids were either transferred directly to the microscope cryoholder or stored in liquid nitrogen. All grids were glow-discharged before use. The micrographs were automatically collected using EPU program (FEI) with a 4 K × 4 K Ceta CMOS digital camera (Gatan, USA).

### 2.5 Extraction, quantification, and western blotting analysis of tissue and AT-EV proteins

The whole lysates of AT or AT-EVs was prepared on ice by using the RIPA lysis buffer (Beyotime, China) containing protease and phosphatase inhibitors (Roche, Germany). After lysis for 20 min, the samples were centrifuged at 12,000 g for 20 min to harvest the supernatant protein solution. Proteins were quantified using a BCA Protein Assay Kit (TIANGEN, China). Western blotting analysis was performed according to previous studies [28, 31]. Briefly, equal amounts of protein samples were loaded onto SDS–PAGE gels and transferred to polyvinylidene fluoride (PVDF) membranes (Millipore, USA) which were blocked with 5% bovine serum albumin (BSA) (Sigma-Aldrich, USA) in TBS at room temperature for 2 h. Then, the membranes were incubated overnight at 4□ with the following primary antibodies: anti-CD9 (ET1601-9, HUABIO, China; diluted at 1:1000), anti-CD63 (sc5275, Santa Cruz Biotechnology, USA; diluted at 1:1000), anti-CD81 (GTX31381, Gene Tex, USA; diluted at 1:1000), anti-Flotillin1 (18634, Cell Signaling Technology, USA; diluted at 1:1000), anti-Caveolin1 (sc53564, Santa Cruz Biotechnology, USA; diluted at 1:1000), anti-Golgin84 (NBP1-83352, Novus Biologicals, USA; diluted at 1:1000), anti-Mitofilin (ab110329, Abcam, UK; diluted at 1:1000), and anti-UCP1 (ab234430, Abcam, UK; diluted at 1:1000). After being washed twice with TBS containing 0.1% Tween-20, the membranes were incubated with peroxidase-conjugated secondary antibodies (Jackson ImmunoResearch, USA) for 1 h at room temperature. The protein bands were visualized by using a chemiluminescence kit (Amersham Biosciences, USA) and captured by a gel imaging system (4600, Tanon, China).

### 2.6 Proteomic analysis

Protein lysates of fresh-isolated AT-EVs were prepared and subjected to LC-MS/MS analysis on a timsTOF Pro mass spectrometry (Bruker Daltonics, USA). The resulting MS/MS data were processed using the MaxQuant search engine (v.1.6.15.0). Proteins were identified by comparing against the Uniport database with a false discovery rate set at 0.01 for both peptides and proteins. The principal component analysis (PCA) and Venn diagram analysis were conducted for the expressed proteins of the three kinds of AT-EVs, and then Mfuzz analysis was performed for the commonly expressed proteins. Next, Venn diagram analysis was conducted for the co-upregulated proteins in BeEVs and BEVs over WEVs (Fold change > 1.5 and *P* value < 0.05), with the top 50 proteins being listed in the heatmap. Moreover, the co-upregulated proteins were included for further functional analysis based on Gene Ontology (GO) and Kyoto Encyclopedia of Genes and Genomes (KEGG) databases.

### 2.7 Biodistribution of AT-EVs *in vivo*

To track the biodistribution of AT-EVs *in vivo*, a fluorescent lipophilic tracer DiR (Invitrogen, USA) was used to label the purified AT-EVs. C57BL/6J mice were injected with labeled AT-EVs *via* the tail vein and then examined under anesthesia by the IVIS Spectrum in vivo imaging system (PerkinElmer, USA) at 24 h post-injection. Moreover, the heart, lung, liver, spleen, kidney, bone, colon, small intestine, BAT, subcutaneous WAT, and visceral WAT were harvested and observed. The fluorescence images were captured and the intensity of fluorescence was quantified with Living Image software (PerkinElmer, USA) [28].

### 2.8 IF staining

Fresh AT samples were fixed in 4% PFA (Biosharp, China) at 4□ for 4 h, washed twice with PBS and dehydrated with 30% sucrose for 16-18 h. After being embedded in optimal cutting temperature (OCT) compound (Leica, Germany), 20 μm cryosections were produced with a cryastat (CM1950, Leica, Germany) and underwent IF staining, according to our previous studies [31, 32]. Briefly, air-dried cryosections were permeabilized by 0.2% Triton X-100 (Sigma-Aldrich, USA) for 20 min at room temperature, blocked in goat serum (Sigma-Aldrich, USA) for 30 min at room temperature and incubated with the primary antibody against mouse UCP1 (ab234430, Abcam, UK; diluted at 1:500) at 4°C for 12 h. Then, the cryosections were washed for three times with PBS and incubated with the Alexa Fluor 594-conjugated secondary antibody (33212ES60, Yeasen Biotechnology, CN; diluted at 1:200) at room temperature for 1 h. Subsequently, after washing with PBS for three times, the slides were mounted with Mounting Medium With DAPI-Aqueous, Fluoroshield (Abcam, UK). Photographs were taken by a confocal microscope (FV1000, Olympus, Japan) and analyzed using the ImageJ software (NIH, USA).

### 2.9 Histological analysis

At sacrifice, fresh dissected tissue samples were fixed in 4% PFA at 4□ for 24 h and underwent dehydration with graded ethanol. Then, 5 μm paraffin-embedded sections were prepared (RM2125, Leica, Germany) and underwent H&E staining [28]. For oil red O staining, dissected liver tissues were flash-frozen with liquid nitrogen and imbedded in OCT compound (Leica, Germany). Then, 15 μm cryo-sections were prepared (CM1950, Leica, Germany), immersed in oil red O working solution for 5 min and rinsed with H_2_O. Sections were counterstained with Mayer’s haematoxylin and mounted. Images were taken using a microscope (M205FA, Leica, Germany) [28].

### 2.10 Lipid contents measurement

For liver tissue sample examination, at sacrifice, the liver tissue of mice was dissected and homogenized in ethyl alcohol buffer on ice according to the manufacturer’s instructions. For blood sample examination, mice were anaesthetized, the whole peripheral blood were obtained from the retro-orbital venous plexus, and the serum was isolated *via* centrifuging at 3,000 rpm for 10 min, as previously described [28]. The TG and total cholesterol (TC) levels in the liver or blood samples were measured by the commercial kits according to the protocols (Nanjing Jiancheng Biology Engineering Institute, China).

### 2.11 RNA isolation and qRT-PCR

Total RNA was extracted from the liver tissue by using Trizol Reagent (Takara, Japan), and complementary DNA (cDNA) was generated using a PrimeScriptTM RT Reagent Kit (Takara, Japan). Then, qRT-PCR was performed with a SYBR Premix Ex Taq II Kit (Takara, Japan) by a Real-Time System (CFX96, Bio-Rad, USA). The relative expression level of each gene was determined after normalization to β*-actin* using the 2^−ΔΔCT^ method, as previously described [28]. Values were expressed as fold changes. All the primer sequences were listed as follows: β*-actin* (Forward: 5’-AACAGTCCGCCTAGAAGCAC-3’; Reverse: 5’-CGTTGACATCCGTAAAGACC-3’); fatty acid synthase (*Fasn*) (Forward: 5’-GGCCCCTCTGTTAATTGGCT-3’; Reverse: 5’-GGATCTCAGGGTTGGGGTTG-3’); sterol regulatory element-binding protein 1 (*Srebf1*) (Forward: 5’-ATGCGGCTGTTGTCTACCAT-3’; Reverse: 5’-GATAGCATCTCCTGCGCACT-3’).

### 2.12 Statistic analysis and schematics

All the data were presented as mean ± standard deviation (SD). Statistical and graph analysis was performed using GraphPad Prism 9.5.1 (GraphPad Software, USA). For multiple group comparisons, significance was assessed by One-way ANOVA with Tukey’s *post hoc* test. Values of *P* < 0.05 were considered statistically significant. All schematics were drawn using Biorender with permission.

## 3 Results

### 3.1 Cold stimulation induces the formation of BeAT

The formation of BeAT was induced by the commonly used method of cold stimulation. In specific, C57BL/6J male mice were subjected to cold (4□) for 3 d, 7 d and 14 d, and the formation of BeAT was evaluated by gross observation and morphological analysis. In line with previous studies [33, 34], with prolonged cold stimulation, the appearance of the inguinal subcutaneous WAT gradually turned into a BAT-like phenotype, which stabilized after 7 days (Figure 1A, Supplemental Figure 1A and 1B). H&E staining also showed that BeAT adipocytes had decreased individual cell size and possessed smaller multilocular lipid droplets compared to white adipocytes which contained a large single lipid droplet, indicating morphological transformation toward brown-like adipocytes (Figure 1B, Supplemental Figure 1C). It has been reported that UCP1, an inner mitochondrial membrane protein, is a hallmark of BAT which mediates the non-shivering thermogenesis of BAT *via* uncoupling mitochondrial respiration from ATP production [35, 36]. To further evaluate the efficiency of cold-induced formation of BeAT, we tested the expression level of UCP1 in the BeAT. IF staining results showed that UCP1 was detected in the BAT and the BeAT after 7 days of cold stimulation, whereas no positive expression was detected in the WAT (Figure 1C). Quantification analysis also revealed that both the average cell area of adipocytes and the expression level of UCP1 in the BeAT were between those of the BAT and the WAT (Figure 1D, E). Moreover, given that the content of BeAT was much greater than that of the BAT (Figure 1F), the BeAT is endowed with a huge advantage in terms of isolation and application. Based on these results, we have successfully established a cold-inducing strategy to efficiently convert WAT into BeAT, which provides a stable and abundant source for isolation of BeEVs.

**Figure 1.**
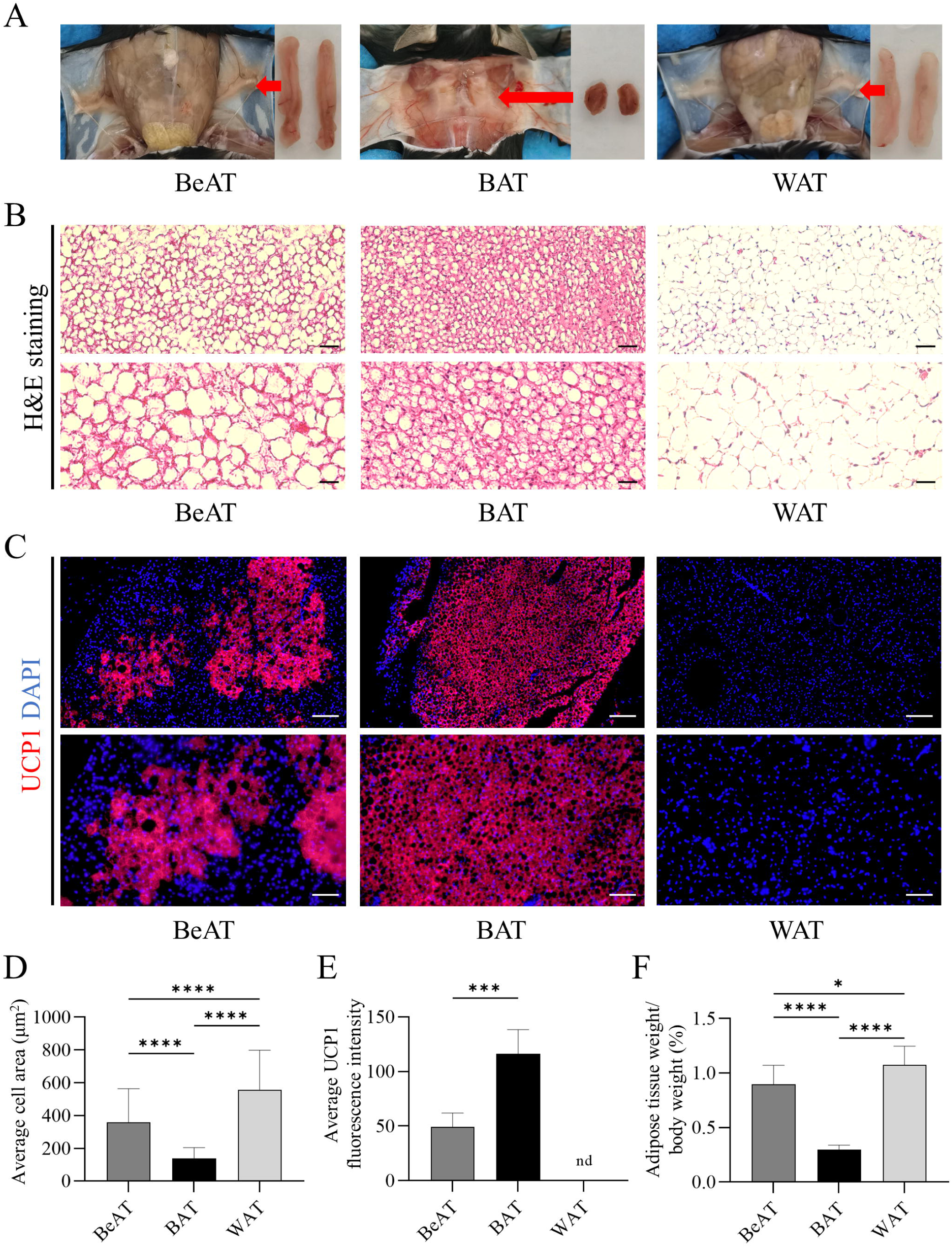
Cold stimulation induces the formation of beige adipose tissue (BeAT). (A) Gross view of the BeAT, brown adipose tissue (BAT) and white adipose tissue (WAT) *in vivo* and after extraction. (B) Representative haematoxylin and eosin (H&E) staining images of the BeAT, BAT and WAT. Scale bars, 50 μm (top) and 25 μm (bottom). (C) Representative immunofluorescent (IF) staining images of uncoupling protein 1 (UCP1) (red) in the BeAT, BAT and WAT counterstained by DAPI (blue). Scale bars, 200 μm (top) and 100 μm (bottom). (D) Quantification of the average cell area of the BeAT, BAT and WAT. (E) Quantification of the average UCP1 fluorescence intensity of the BeAT, BAT and WAT. (F) Quantification of the percentage of adipose tissue weight in body weight of the BeAT, BAT and WAT. Data are presented as mean ± SD. Statistical analyses are performed by One-way ANOVA with Tukey’s post hoc test. *, *P* < 0.05; ***, *P* < 0.001; ****, *P* < 0.0001; nd, not detected.

### 3.2 BeEVs are isolated with high yield and typical EV characteristics

For now, the standard isolating method for tissue-derived EVs remains controversial [9]. The conventional methods mainly involve harvest from the tissue culture medium (which needs more than 24 hours) or release from the extracellular space by direct digestion of tissue with collagenase I and IV (which needed approximately 6 hours) [8]. In this field, our group has recently developed and optimized an enzyme digestion and differential centrifugation-based protocol to isolate EVs from tissues [30]. Using this method, we have isolated EVs from the three kinds of AT (Figure 2A) and characterized them according to the Minimal information for studies of extracellular vesicles 2018 (MISEV2018) [37]. NTA measurement showed that the diameters of BeEVs ranged from 100 to 800 nm and peaked at 150-250 nm, which was consistent with those of WEVs and BEVs (Figure 2B). Morphological observation through TEM and, more precisely, through Cryo-EM showed that BeEVs exhibited a typical sphere-like morphology with peripheral bilayer membrane, similar to BEVs and WEVs (Figure 2C, D). Moreover, western blotting analysis revealed that all the three types of AT-EVs and their origin AT expressed certain EV markers, including CD9, CD63, Flotillin1, and mitofilin, while CD81 and Golgin84 were only expressed in the AT but not in the AT-EVs (Figure 2E). Of note, both BeEVs and BEVs as well as their source tissue expressed UCP1 while WEVs and WAT showed no expression (Figure 2E), consistent with the above IF results. More importantly, statistical analysis showed that although with similar diameters, the particle number of BeEVs harvested from the BeAT is significantly higher than that of WEVs derived from the WAT of equal mass (Figure 2F, G). We further calculated the proportion of EV protein within the total AT protein and found that BeEVs held a significantly higher proportion than WEVs (Figure 2H). Taken together, these data indicate that BeEVs possess the typical characteristics of EVs in size, morphology, structure, and protein markers, and own the advantages of abundant source, easy access and high yield.

**Figure 2.**
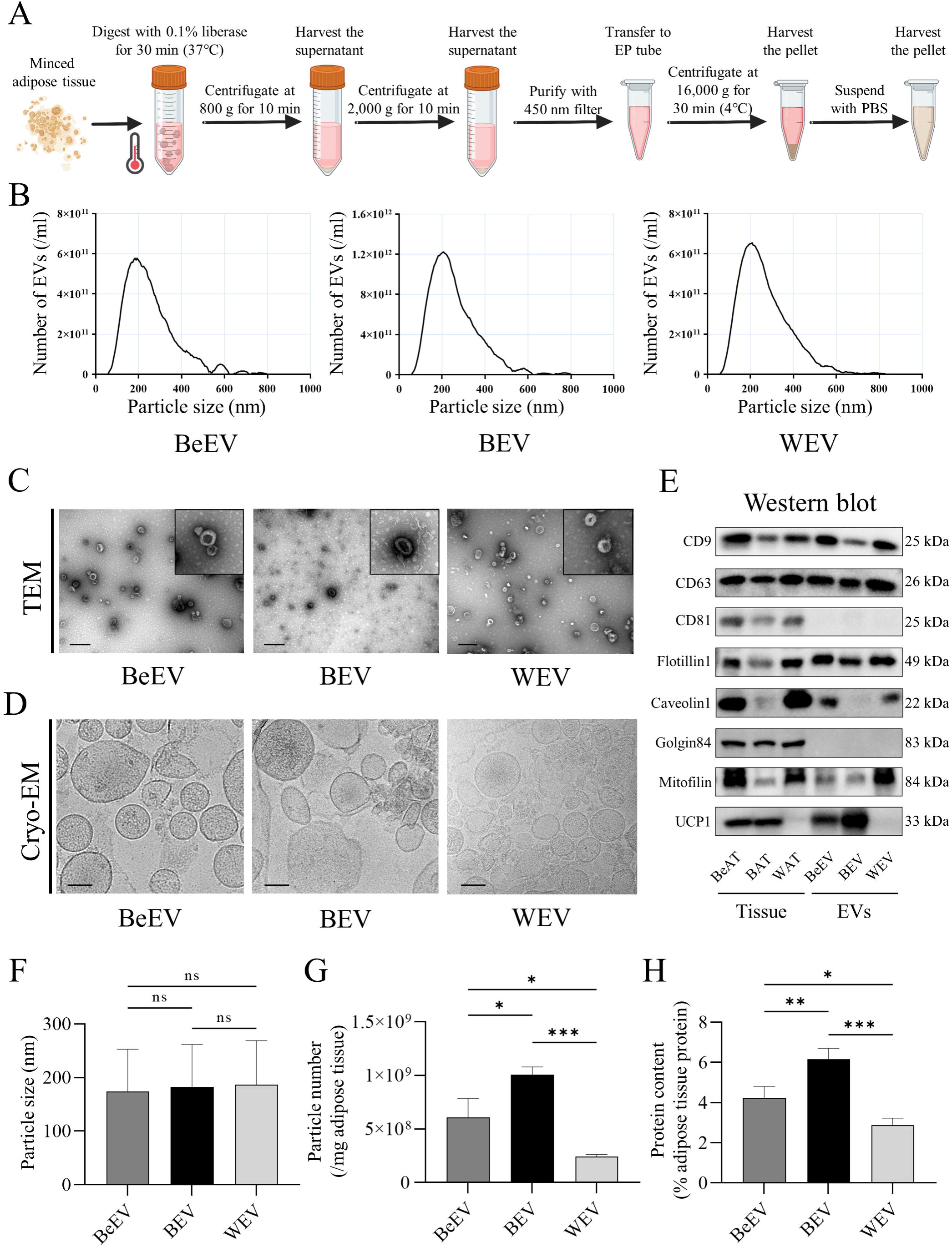
BeAT-derived extracellular vesicles (BeEVs) are isolated with high yield and typical EV characteristics. (A) Schematic diagram showing the isolation procedure of AT-derived EVs. (B) Nanoparticle tracking analysis (NTA) showing the particle number and the size distribution of BeEVs, BAT-derived EVs (BEVs) and WAT-derived EVs (WEVs). (C and D) Representative transmission electron microscope (TEM) and cryo-electron microscopy (Cryo-EM) images showing the morphology of BeEVs, BEVs and WEVs. Scale bar, 500 nm (TEM) and 100 nm (Cryo-EM). (E) Western blotting analysis of the protein markers of the BeEVs, BEVs and WEVs, as well as the BeAT, BAT and WAT. (F) Quantification of the particle size of the BeEVs, BEVs and WEVs. (G) Quantification of the particle number of the BeEVs, BEVs and WEVs. (H) Quantification of the percentage of AT-EV protein content in total AT protein. Data are presented as mean ± SD. Statistical analyses are performed by One-way ANOVA with Tukey’s post hoc test. *, *P* < 0.05; **, *P* < 0.01; ***, *P* < 0.001; ns, *P* > 0.05.

### 3.3 BeEVs possess distinct protein profiles with functional implications

The cargo composition, especially the contained proteins, is key in determining the functions of EVs [28]. To comprehensively analyze the protein composition of BeEVs, we prepared protein samples of the three kinds of AT-EVs and performed proteomic analysis of BeEVs, with WEVs and BEVs as references. A total of 5026 proteins were identified, of which 3803 were quantified and used for further analysis. PCA analysis showed that the protein composition of BeEVs was significantly different from that of BEVs and WEVs, indicating a distinct protein profile of BeEVs (Figure 3A). We further performed Venn analysis of the proteins expressed in each group and found that a total of 2631 proteins were co-expressed in the three kinds of AT-EVs, while 32, 47, 282 proteins were solely expressed by BeEVs, BEVs and WEVs, respectively (Figure 3B). In order to further explore the expression patterns of the commonly expressed 2631 proteins, we performed Mfuzz analysis. Among the 6 patterns (clusters 1-6), proteins of BeEVs and WEVs showed similarity in clusters 1 and 3, which were mainly located in the membrane and were related to “Endocytosis”, “Protein binding”, and “Vesicle-mediated transport” (Figure 3C). Meanwhile, proteins with similar expression patterns of BeEVs and BEVs (cluster 5 and 6) are mainly enriched in the mitochondrion, and are associated with “Thermogenesis”, “NAFLD”, “Mitochondrion organization”, “Cellular respiration”, and “biosynthesis of unsaturated fatty acids” (Figure 3C). These results suggest that BeEVs have a distinctive protein composition with functional implications.

**Figure 3.**
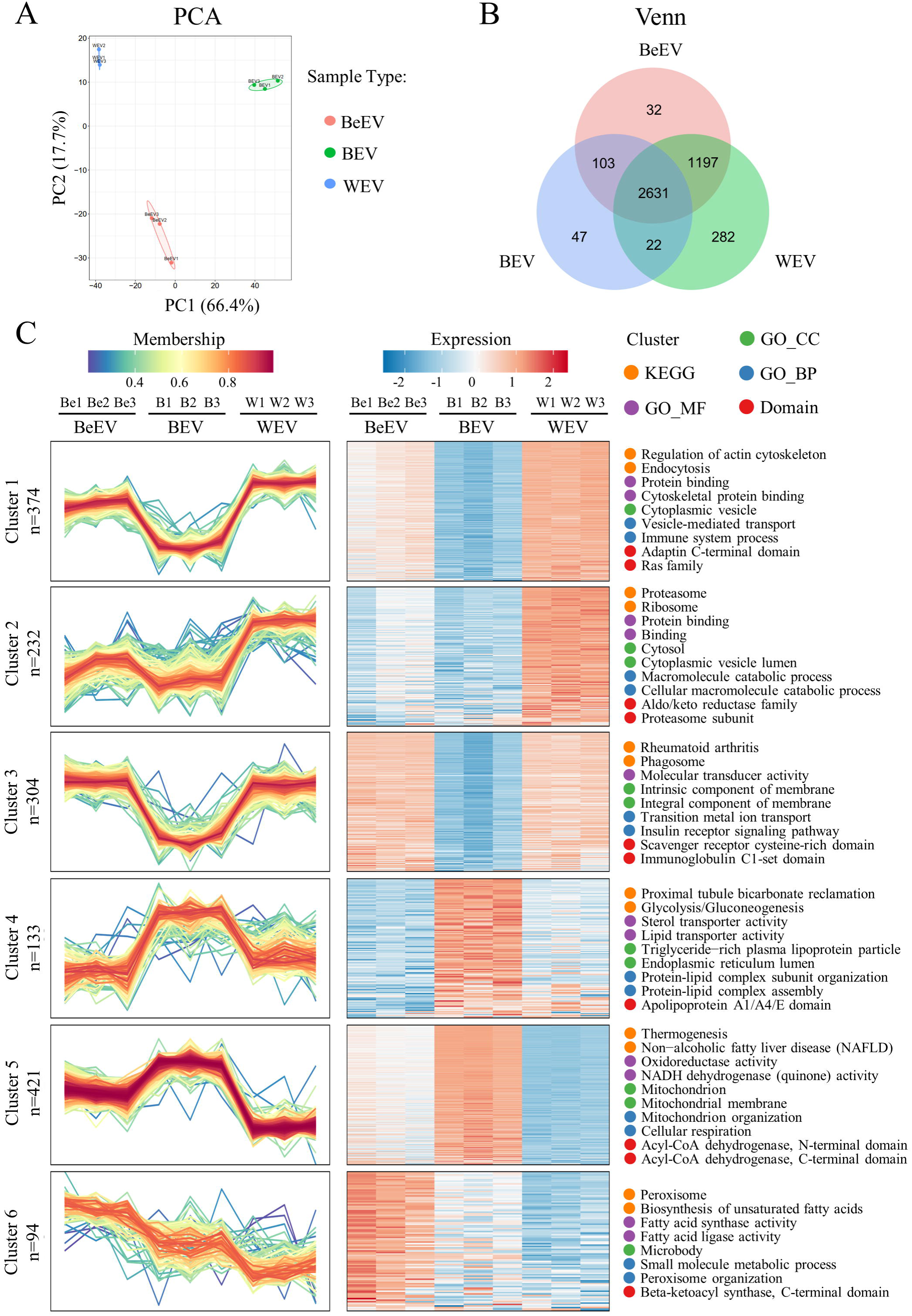
Proteomics analysis shows that BeEVs possess distinct protein profiles with functional implications. (A) Principal component analysis (PCA) analysis of the expressed proteins of BeEVs (orange dots and ellipse), BEVs (green dots and ellipse) and WEVs (blue dots and ellipse). (B) Venn diagram showing the specific number of solely expressed or co-expressed proteins in the BeEVs (orange circle), BEVs (blue circle) and WEVs (green circle). (C) Mfuzz analysis showing the expression patterns of the commonly expressed proteins in the BeEVs, BEVs and WEVs. Each cluster is displayed by a line graph and a heatmap. Line chart: the X-axis represents the samples, the Y-axis represents the relative protein expression, a broken line represents a protein, and the color of the line represents the membership of the protein within the cluster. Heat map: the X-axis represents the samples, the Y-axis represents different proteins, and the color of each box represents the relative expression of proteins in samples. Spot color: orange for Kyoto Encyclopedia of Genes and Genomes (KEGG); purple for Gene Ontology_molecular function (GO_MF); green for Gene Ontology_Cellular Component (GO_CC); blue for Gene Ontology_Biological Process (GO_BP); red for Domain.

### 3.4 BeEVs are enriched with metabolic regulatory proteins

Given that BeAT is known as browning WAT that possesses BAT-like functions in the regulation of energy metabolism, we further focused on the proteins of BeEVs which might mediate the functions. We first performed Venn analysis of the proteins which were upregulated in BeEVs and BEVs, respectively over WEVs (Figure 4A). Results showed that 423 proteins were commonly upregulated in BeEVs and BEVs over WEVs, while 125 and 174 proteins were separately upregulated in BeEVs and BEVs over WEVs, supporting functional similarity between BeEVs and BEVs (Figure 4A). As shown by the heatmap, the top 50 proteins of the 423 co-upregulated proteins mainly included the UCP family responsible for energy expenditure (UCP1, UCP3) [36, 37], the solute carriers (SLC) family of transmembrane transport proteins (Slc27a2, Slc25a42, Slc25a20, Slc25a35, Slc25a16) [38, 39], the mitochondrial ribosomal protein (MRP) family involved in mitochondrial ribosomal composition (Mrpl38, Mrpl4, Mrpl46, Mrpl47) [40], and the cytochrome c oxidase (COX) family essential for the mitochondrial respiratory chain (Cox7a1, Cox7b, Cox6a1) [41] (Figure 4B). These results showed that BeEVs were enriched with a large number of proteins involved in energy metabolism and transmembrane transport. We further conducted functional analysis among these 423 co-upregulated proteins. According to the GO enrichment analysis, these proteins are predominantly located in the mitochondria within “Cellular component (CC)” category (Figure 4C) and highly involved in metabolic processes, including “generation of precursor metabolites and energy”, “oxidative phosphorylation”, “fatty acid beta-oxidation” and “mitochondrial transmembrane transport” within the “Biological process (BP)” category (Figure 4D), as well as “acyl-CoA dehydrogenase activity”, “electron transfer activity”, “oxidative phosphorylation uncoupler activity” and “transmembrane transporter activity” within the “Molecular function (MF)” category (Figure 4E). KEGG enrichment analysis also revealed that these proteins are closely associated with the metabolic-related pathways, such as “Citrate cycle (TCA cycle)”, “Fatty acid degradation”, “Oxidative phosphorylation”, “PPAR signaling pathway”, “Thermogenesis”, and “NAFLD-related pathway” (Figure 4F). Collectively, these results strongly indicate that BeEVs possess a protein basis for performing BEV-like metabolic regulation functions, which endows it with potential therapeutic effects on metabolic diseases.

**Figure 4.**
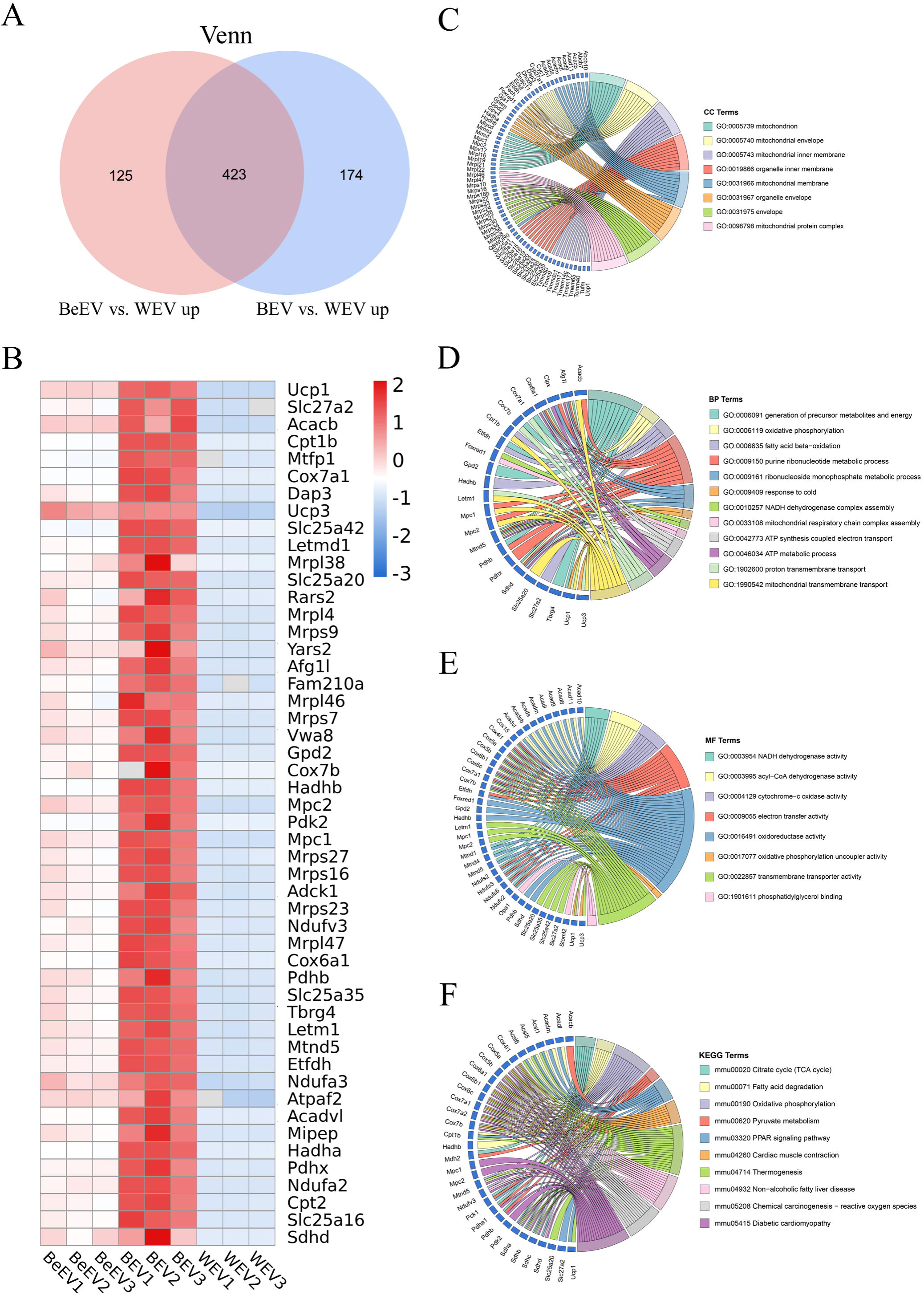
BeEVs are enriched with metabolic regulatory proteins. (A) Venn diagram showing the number of co-upregulated proteins in BeEVs and BEVs over WEVs (Fold change > 1.5 and *P* value < 0.05). (B) Heatmap showing the top 50 proteins of the co-upregulated proteins in BeEVs and BEVs over WEVs. (C) Chord diagram of GO enrichment analysis of the co-upregulated proteins in BeEVs and BEVs over WEVs, within the “Cellular component (CC)” category. (D) Chord diagram of GO enrichment analysis of the co-upregulated proteins in BeEVs and BEVs over WEVs, within the “Biological process (BP)” category. (E) Chord diagram of GO enrichment analysis of the co-upregulated proteins in BeEVs and BEVs over WEVs, within the “Molecular function (MF)” category. (F) Chord diagram of KEGG enrichment analysis of the co-upregulated proteins in BeEVs and BEVs over WEVs.

### 3.5 Systematically injected BeEVs engraft into the liver and alleviate HFD-induced metabolic dysfunction

Based on the above observation, we proceeded to investigate the therapeutic potential of BeEVs. First, we detected the biodistribution of systemically delivered BeEVs in various organs (BAT, subcutaneous WAT, visceral WAT, heart, lung, liver, spleen, kidney, bone, colon, small intestine) and found significant accumulation of BeEVs in the liver at 24 h post-injection (Figure 5A), which was consistent with previous studies showing hepatic tropism for exogenous nanoparticles [42]. Given that the liver is a center organ in body metabolism, we next validated the therapeutic efficiency of BeEVs in a HFD-induced model of NAFLD, a leading chronic liver disease worldwide [43]. In specific, a sequential injection protocol of BeEVs with an interval of 1 week was conducted for 8 weeks (Supplemental Figure 2). As expected, compared to control mice, HFD feeding led to an obese phenotype in mice which was relieved in both the HFD+PBS and HFD+WEV groups but not in the HFD+WEV group (Figure 5B). Consistently, biochemical assays showed that exogenous injection of BeEVs and BEVs both significantly reduced fasting glucose and TC levels in serum, compared with the HFD+PBS group, whereas WEVs had no beneficial effects on these indices (Figure 5C-E). In addition, quantification analysis found that the HFD+BeEV and HFD+BEV groups had significantly lower body weight and weight gain than the HFD+PBS group, while the HFD+WEV group showed no significant difference from the HFD+PBS group (Figure 5F, G). Moreover, the liver weight of the HFD+BeEV and HFD+BEV groups was remarkably reduced compared with the HFD+PBS group while no difference was detected between the HFD+PBS and HFD+WEV groups (Figure 5H). Accordingly, these data exhibited that the exogenously injected BeEVs mainly engrafted into the liver and effectively alleviated HFD-induced metabolic dysfunction.

**Figure 5.**
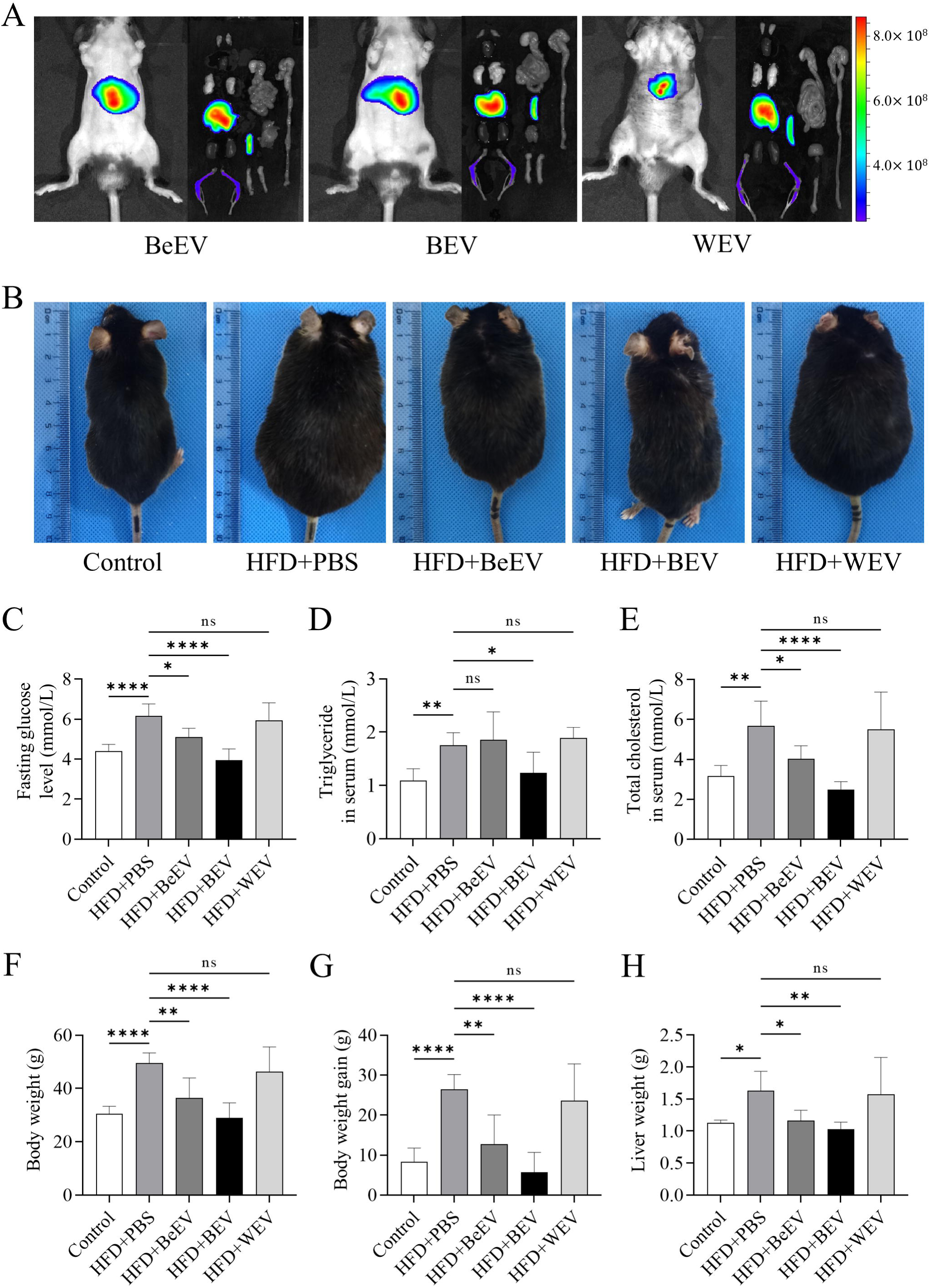
Systematically injected BeEVs engraft into the liver and alleviate high fat diet (HFD)-induced metabolic dysfunction. (A) Biodistribution of exogenously injected DiR-labelled BeEVs *in vivo* and *ex vivo*. (B) Photographs of C57BL/6J mice of indicated groups. HFD+PBS, mice feeding HFD and treated with PBS; HFD+BeEV, mice feeding HFD and treated with BeEVs; HFD+BEV, mice feeding HFD and treated with BEVs; HFD+WEV, mice feeding HFD and treated with WEVs. Each small grid of the background represents 1 mm. (C) Quantification of the fasting glucose level. (D) Quantification of the level of triglyceride in serum. (E) Quantification of the level of total cholesterol in serum. (F) Quantification of the body weight. (G) Quantification of the body weight gain. (H) Quantification of the liver weight. Data are presented as mean ± SD. Statistical analyses are performed by One-way ANOVA with Tukey’s post hoc test. *, *P* < 0.05; **, *P* < 0.01; ****, *P* < 0.0001; ns, *P* > 0.05.

### 3.6 BeEV infusion rescues liver steatosis *via* reducing lipogenesis

The above findings prompt us to further explore the direct effects of BeEVs on fatty liver. By comparing the dissected livers, we found hypertrophic and steatosis appearance in the HFD+PBS and HFD+WEV groups, while the HFD+BeEV and HFD+BEV groups showed normal size and appearance (Figure 6A). H&E staining provided further evidence that the vacuole-like structures diffused in the liver tissues in the HFD+PBS and HFD+WEV groups, associated with steatosis ballooning in hepatocytes. In contrast, these pathologic changes scarcely appeared in the HFD+BeEV group, and the HFD+BEV and control groups showed normal morphologies (Figure 6B). Similarly, oil red O staining and relevant quantification also showed that the area of lipid deposition was significantly decreased in the HFD+BeEV and HFD+BEV groups, compared with the HFD+PBS and HFD+WEV groups (Figure 6C, D).

**Figure 6.**
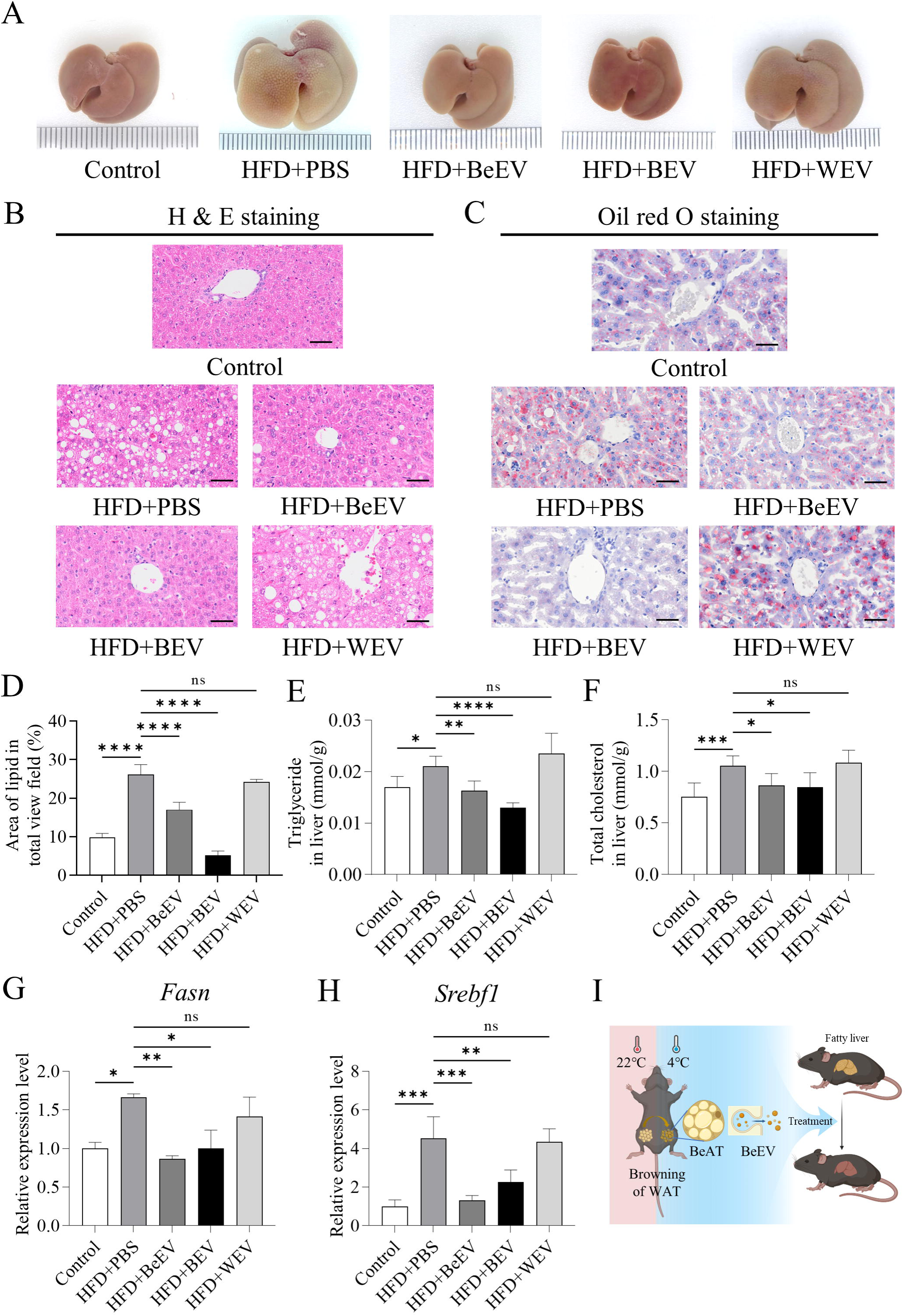
BeEV infusion rescues liver steatosis *via* reducing lipogenesis. (A) Gross view of the livers extracted from the mice of indicated groups. (B) Representative H&E staining images of liver tissues. Scale bars, 50 μm. (C and D) Representative oil red O staining images of liver tissues and the corresponding quantification of the percentage of aera of lipid in total view field. Scale bars, 25 μm. (E) Quantification of the level of triglyceride in liver. (F) Quantification of the level of total cholesterol in liver. (G and H) Quantitative real time polymerase chain reaction (qRT-PCR) analysis of the mRNA expression levels of fatty acid synthase (*Fasn*) (G) and sterol regulatory element-binding protein 1 (*Srebf1*) (H) in liver tissue, normalized to β*-actin*, and quantification of fold changes over the Control group. (I) Schematic diagram showing the synopsis of the findings. Data are presented as mean ± SD. Statistical analyses are performed by One-way ANOVA with Tukey’s post hoc test. *, *P* < 0.05; **, *P* < 0.01; ***, *P* < 0.001; ****, *P* < 0.0001; ns, *P* > 0.05.

As for biochemical assays of liver lysates, levels of TG and TC were significantly lower in the HFD+BeEV and HFD+BEV groups than in the HFD+PBS group, whereas there were no significant differences between the HFD+WEV and HFD+PBS groups (Figure 6E, F). Based on the histological and biochemical results, we further conducted qRT-PCR experiment to analyze the lipogenesis activity in the liver. Results showed that the levels of key lipogenesis genes (*Fasn* and *Srebf1*) in liver were remarkably decreased by systemic injection of BeEVs and BEVs, while WEVs exerted no such effects (Figure 6G, H), suggesting that BeEVs alleviate liver steatosis *via* inhibiting the synthesis of lipids. Last but not least, the safety of exogenously injected BeEVs was tested so as to promote their translational application. As shown by H&E staining and quantification of weight, after AT-EV injection, the major organs of the individuals, such as the heart, spleen, lung, and kidney, remain normal histomorphology and weight (Supplemental Figures 3). Taken together, the main *in vivo* findings are summarized as follows: cold stimulation induces formation of BeAT which produce BeEVs that are enriched with functional proteins involved in metabolic regulation and provide a promising strategy for NAFLD therapy (Figure 6I).

## 4 Discussion

Since EVs were firstly introduced in the 1980s [44], this field has been focused on cell culture-derived and body fluid-derived EVs, while the studies on tissue-derived EVs are still limited [10]. Of note, within recent years, tissue-derived EVs are garnering increasing attention, which possess plentiful biological information inherited from the parental tissue, and are equipped with easy access and high yield [8]. As a central metabolic organ, AT is widely distributed throughout the body and engages in crosstalk with multiple organs, which is an important source of EVs *in vivo* [8]. While the EVs of the classical WAT and BAT have been investigated by a few studies [11–13], much less is known about the EVs of BeAT, a thermogenic AT induced from WAT. The present study for the first time isolates and characterizes BeEVs, identifies their protein profile and more importantly, reveals that BeEVs, enriched with mitochondrial-related functional proteins, can effectively alleviate NAFLD.

In the last decade, WAT browning has become a hotspot of research, with high potential as a novel therapeutic strategy for obesity and metabolic diseases [45, 46]. Of current studies, the main methods used to induce BeAT formation include cold exposure, starvation, hormones, exercise, and hormones [21, 47], among which cold stimulation is the most commonly used method. It is worth noting that BeAT is not stable after *in vivo* induction and can be converted back to WAT without the stimulation factors [48]. Therefore, the investigation and application of BeAT-derived EVs, which are recognized to preserve components and functions of source cells, are likely to promote the translational application of BAT. In the present study, we established a fast and efficient method for isolating BeEVs based on our previously established protocol [30]. Of note, a key step is the use of Liberase for tissue digestion, a purified enzyme blend with mild action, high efficiency, and good biosecurity for isolating EVs from tissue. This is the first report for the isolation of BeEVs and our results show that the extracted BeEVs possess the typical features of EVs, thus promoting the functional explorations of BeEVs and providing reference for the follow-up studies. Moreover, considering that BeEVs are derived from different cell sources and the inherent heterogeneity of EVs from one kind of cell, further investigations are needed so as to further deepen the understanding of BeEVs.

Latest research shows that the assembly of EV cargo is an active and selective process, rather than a result of random loading [49]. Thus, the molecular composition of EVs is a key focus in the field, as they represent the source of EVs and fulfill the functions of EVs. Of note, advances in high-throughput sequencing have provided an effective tool to analyze the composition of AT-EVs [50]. For example, BAT-derived exosomes are enriched with mitochondria components, compared with serum-derived exosomes [12]. Besides, the EVs of WAT and BAT isolated from lean or overweight individuals show difference in functional proteins and lipids [51, 52]. In this study, we are the first to map the protein expression profile of BeEVs with the proteome maps of BEVs and WEVs as the reference. In terms of total protein profile, BeEVs show both similarities and independence with BEVs and WEVs, while in terms of functional protein expression, BeEVs are enriched with multiple metabolic regulation proteins similar to BEVs. Notably, a variety of mitochondrial-related proteins were enriched in BeEVs, such as UCP1 [35, 36]. We assume that UCP1-dependent energy dissipation contributes to the improvement of lipid metabolism induced by BeEVs, but the specific pathways remain to be revealed. It is also worth noting that there are 32 unique proteins solely expressed in BeEVs, which may endow BeEVs with novel functions that require further exploration.

NAFLD is a worldwide prevalent metabolic disease lacking effective treatments, which is characterized by unbalanced lipogenesis and lipolysis in the liver [23, 24]. EVs, especially stem cell-derived EVs, has been demonstrated as an effective remedy in liver disease treatment but is restricted by the long culture period and low yield [8, 9]. In this study, we have provided a proof-of-concept study that exogenously injected BeEVs could effectively reduce liver lipogenesis and thus alleviate liver steatosis in HFD-feed mice. Considering that BeEVs are easily obtained and stored, with abundant yield and liver-targeting property, our findings will provide a potent rationale for translational paradigms and promote the establishment of optimized AT-derived EV-based therapies. Furthermore, as critical players in delivering messages between different cells and organs, AT-EVs have been studied in both *in vivo* physiological and pathological conditions, such as systemic glucose homoeostasis maintenance, exercise cardioprotection, insulin resistance, diabetic cognitive impairment, and diabetic ischemia/reperfusion Injury [6, 7, 13, 53, 54]. Nevertheless, the roles of endogenous BeEVs requires further study and exploration.

In conclusion, our study extends the current understanding of AT-EVs and opens a new window regarding the isolation, identification, protein composition and functional analysis of BeEVs, a potent metabolic regulator and a novel remedy for metabolic diseases.

## Acknowledgments

This work is supported by grants from the National Natural Science Foundation of China (82171568, 82301028, 82170988, 82371020, 11774279, 11774280 and 82201014), China Postdoctoral Science Foundation (BX20230485), the National Science Fund for Outstanding Young Scholars (11922410), the Young Science and Technology Rising Star Project of Shaanxi Province (2023KJXX-027), Shaanxi Province Innovation Capability Support Program Scientific and Technological Innovation Team (2023-CX-TD-69), the National Defense Biotechnology Outstanding Young Talents Fund (01-SWKJYCJJ24), Shaanxi Province Science Fund For Distinguished Young Scholars (2022JC-52), and the Progressive Project of The Air Force Medical University (2020JSTS11). We would like to sincerely thank Dr. Guorong Deng for his crucial help with the data analysis procedure.

## Author Contributions

Kai Zhang, Sha Zhang and Bing-Dong Sui contributed equally to the experimental performing, data acquisition and analysis, and manuscript drafting. Yuan Yuan and Liu Lu contributed to animal experiments. Si-Qi Ying and Cheng-Han Li contributed to data analysis and interpretation. Kai-Chao Zhang, Shu-Juan Xing and Jin Liu contributed to histological section staining and analysis. Zhi-Wei Yang contributed to Cyro-EM data acquisition. Yang Sun, Li-Juan Yu and Yan Jin contributed to data interpretation and manuscript revision. Bing-Dong Sui, Chen-Xi Zheng and Liang Kong contributed to the study conception and design, data interpretation and manuscript revision. All authors have read and approved the current version of the manuscript.

## Data Availability Statement

All data presented in this study are available on reasonable request from the corresponding authors.

---

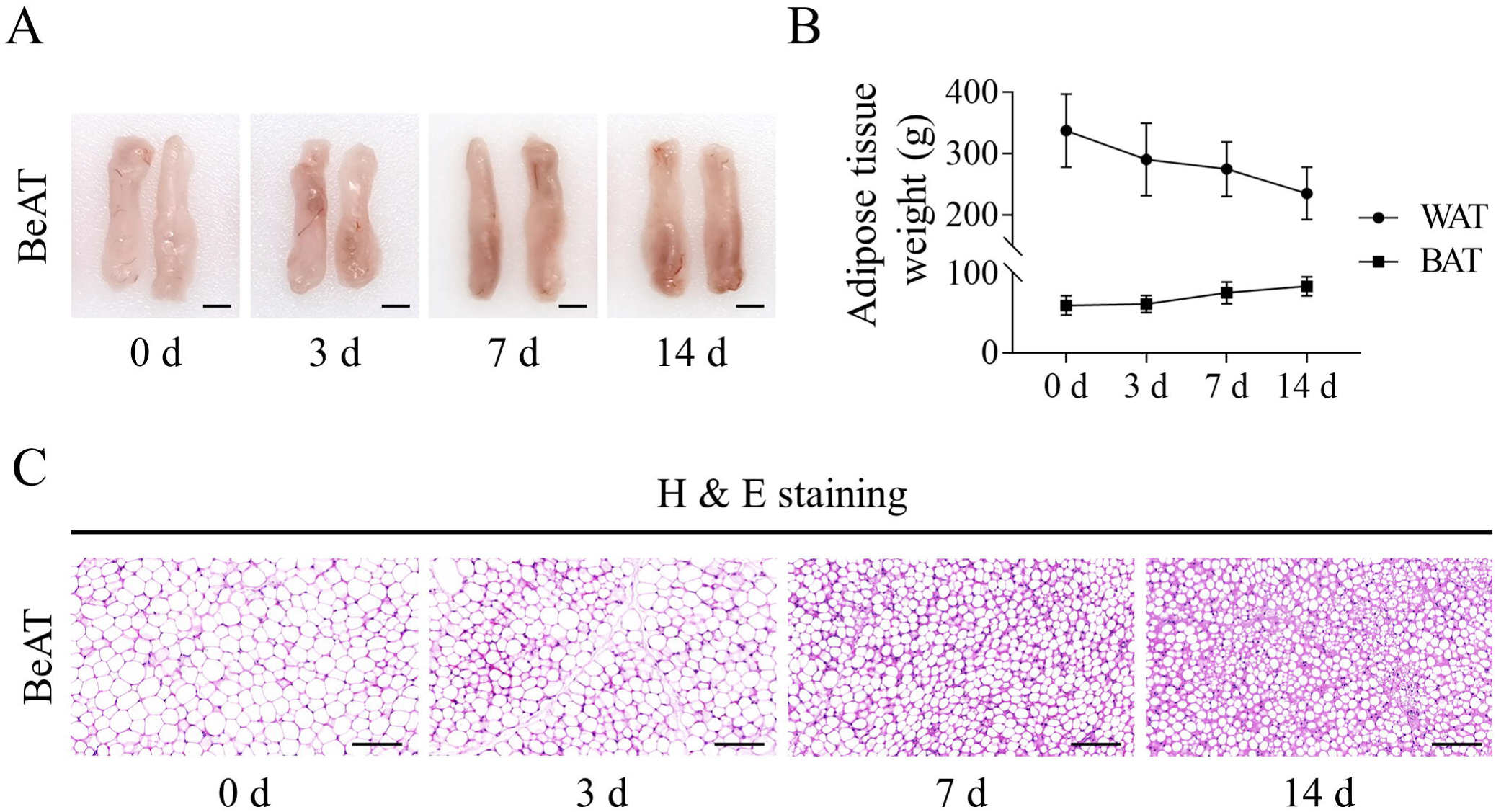

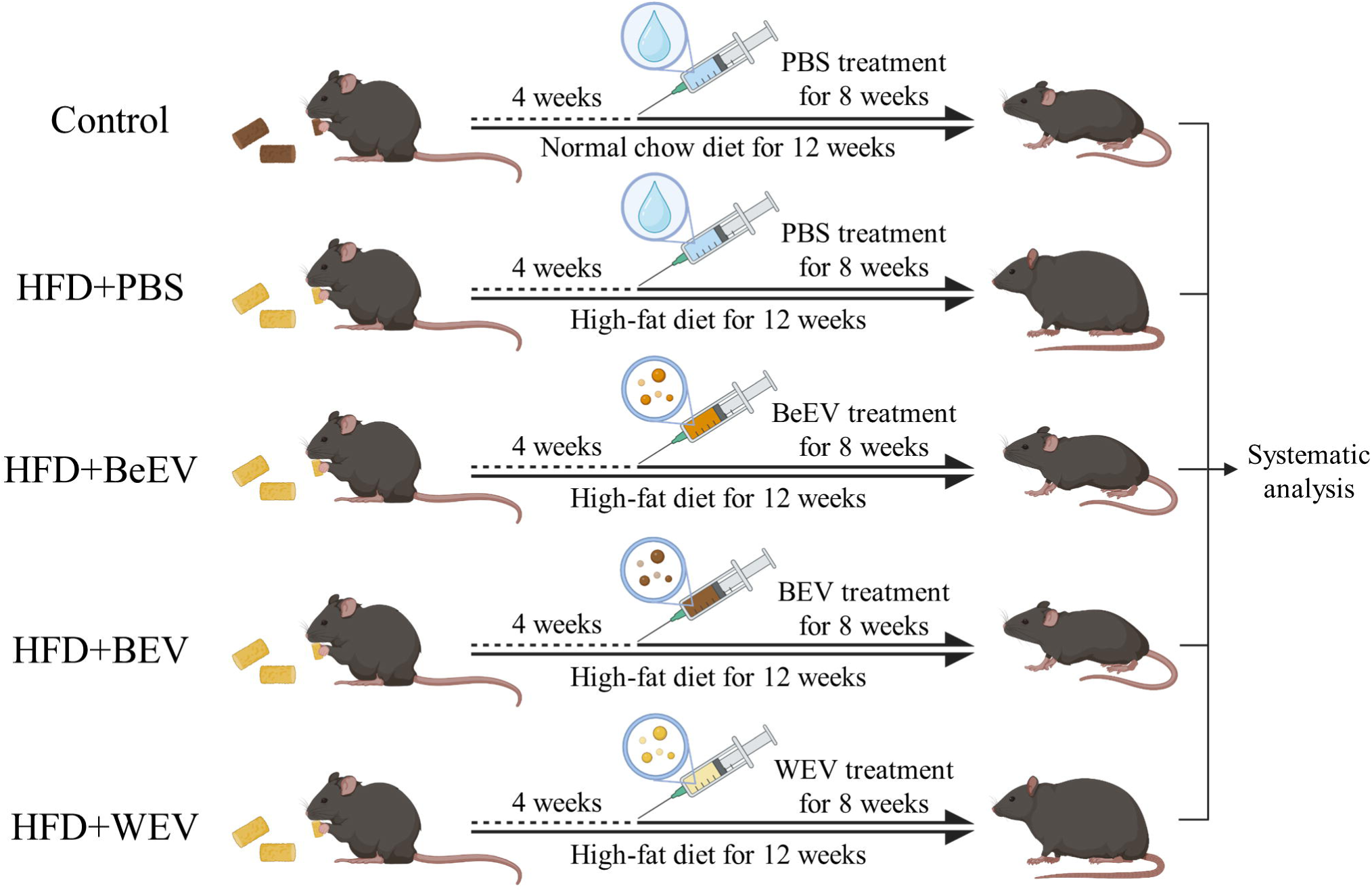

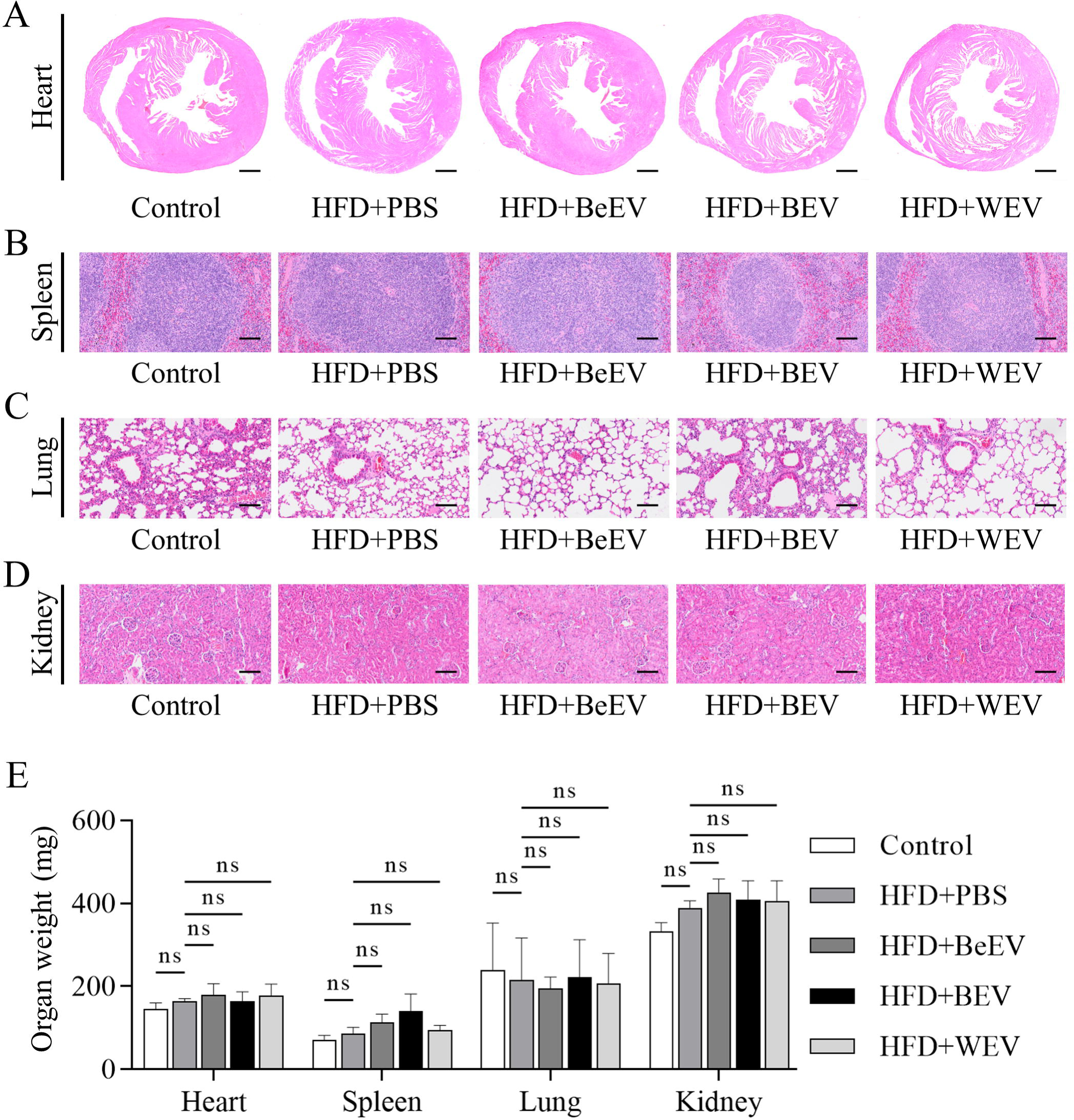

