## Supplemental Figure Legends for "Beige adipose tissue-derived extracellular vesicles: a potent metabolic regulator and a novel remedy for nonalcoholic fatty liver disease"

**Supplementary figures and legendsSupplementary Figure 1. Formation of beige adipose tissue (BeAT) induced by prolonged cold stimulation.**

(A) Gross view of the subcutaneous white adipose tissue (WAT) undergoing browning induced by prolonged cold stimulation. Scale bars, 500 μm. (B) Quantification of the weight of WAT and brown adipose tissue (BAT) with the prolonged cold stimulation. (C) Representative H&E staining images of the subcutaneous WAT with the prolonged cold stimulation. Scale bars, 50 μm.

**Supplementary Figure 2. Establishment of high-fat diet (HFD)-induced nonalcoholic fatty liver disease (NAFLD) model and the application of systemically injected AT-derived extracellular vesicles (AT-EVs).**

Four-week-old male C57BL/6J mice were placed on a HFD or a normal chow diet during the whole process of the experiment. After 4 weeks of HFD feeding, the mice were intravenously administrated with BeAT-derived EVs (BeEVs), BAT-derived EVs (BEVs), WAT-derived EVs (WEVs) (appropriately 100 μg based on protein measurement) or equivalent amount of PBS once a week for 8 weeks. Systematic assays were conducted at the indicated times.

**Supplementary Figure 3. Biosecurity analysis of the exogenously injected BeEVs.**

(A) Representative H&E staining images of the heart tissue in indicated groups. Scale bars, 500 μm. (B) Representative H&E staining images of the spleen tissue in indicated groups. Scale bars,100 μm. (C) Representative H&E staining images of the lung tissue in indicated groups. Scale bars, 100 μm. (D) Representative H&E staining images of the kidney tissue in indicated groups. Scale bars, 100 μm. (E) Quantification of the weight of the heart, spleen, lung and kidney in indicated groups. Data are presented as mean ± SD. Statistical analyses are performed by One-way ANOVA with Tukey’s post hoc test. ns, *P* > 0.05.
